# A new method for detecting autocorrelation of evolutionary rates in large phylogenies

**DOI:** 10.1101/346635

**Authors:** Qiqing Tao, Koichiro Tamura, Fabia Battistuzzi, Sudhir Kumar

## Abstract

New species arise from pre-existing species and inherit similar genomes and environments. This predicts greater similarity of mutation rates and the tempo of molecular evolution between direct ancestors and descendants, resulting in autocorrelation of evolutionary rates within lineages in the tree of life. Surprisingly, molecular sequence data have not confirmed this expectation, possibly because available methods lack power to detect autocorrelated rates. Here we present a machine learning method to detect the presence evolutionary rate autocorrelation in large phylogenies. The new method is computationally efficient and performs better than the available state-of-the-art methods. Application of the new method reveals extensive rate autocorrelation in DNA and amino acid sequence evolution of mammals, birds, insects, metazoans, plants, fungi, and prokaryotes. Therefore, rate autocorrelation is a common phenomenon throughout the tree of life. These findings suggest concordance between molecular and non-molecular evolutionary patterns and will foster unbiased and precise dating of the tree of life.

## Introduction

Rates of molecular sequence evolution vary extensively among species (Ho and Duchêne 2014; Kumar and Hedges 2016; dos Reis et al. 2016). The causes and consequences of this evolutionary rate variation are of fundamental importance in molecular phylogenetics and systematics (Kimura 1983; Lanfear et al. 2010; Lynch 2010), not only to inform about the relationship among molecular, biological, and life history traits, but also as a prerequisite for reliable estimation of divergence times among species and genes (Ho and Duchêne 2014; Kumar and Hedges 2016). Three decades ago, Gillespie (1984) proposed that molecular evolutionary rates within a phylogeny will be autocorrelated due to similarities in genomes, biology and environments between ancestral species and their immediate progeny. This idea led to statistical modelling of the variability of evolutionary rates among branches and formed the basis of the earliest relaxed clock methods for estimating divergence times without assuming a strict molecular clock (Sanderson 1997; Thorne et al. 1998; Kumar 2005; Ho and Duchêne 2014; Kumar and Hedges 2016). However, the independent branch rate (IBR) model has emerged as a strong alternative to the autocorrelated branch rate (ABR) model. IBR posits that rates vary randomly throughout the tree, such that the evolutionary rate similarity between an ancestor and its descendant is, on average, no more than that between more distantly-related branches in a phylogeny (Drummond et al. 2006; Ho and Duchêne 2014).

The IBR model is now widely used in estimating divergence times from molecular data for diverse groups of species, including mammals (Drummond et al. 2006), birds (Brown et al. 2008; Claramunt and Cracraft 2015; Prum et al. 2015), amphibians (Feng et al. 2017), plants (Moore and Donoghue 2007; Bell et al. 2010; Smith et al. 2010; Linder et al. 2011; Lu et al. 2014; Barreda et al. 2015; Barba-Montoya et al. 2018), and viruses (Drummond et al. 2006; Buck et al. 2016; Metsky et al. 2017). If the IBR model best explains the variability of evolutionary rates, then we must infer a decoupling of molecular and biological evolution, because morphology, behavior, and other life history traits are more similar between closely-related species (Sargis and Dagosto 2008; Lanfear et al. 2010; Cox and Hautier 2015) and are correlated with taxonomic or geographic distance (Wyles et al. 1983; Shao et al. 2016).

Alternatively, the widespread use of the IBR model (Drummond et al. 2006; Moore and Donoghue 2007; Brown et al. 2008; Bell et al. 2010; Smith et al. 2010; Linder et al. 2011; Lu et al. 2014; Claramunt and Cracraft 2015; Prum et al. 2015; Buck et al. 2016; Feng et al. 2017; Metsky et al. 2017) may be explained by the fact that the currently available statistical tests lack sufficient power to reject the IBR model (Ho et al. 2015). In fact, some studies report finding extensive branch rate autocorrelation (e.g., Lepage et al. (2007)), but others do not agree (e.g., Linder et al. (2011)). Consequently, many researchers use both ABR and IBR models when applying Bayesian methods to date divergences (Wikström et al. 2001; Drummond et al. 2006; Bell et al. 2010; Erwin et al. 2011; Meredith et al. 2011; dos Reis et al. 2012; Magallón et al. 2013; Jarvis et al. 2014; Hertweck et al. 2015; dos Reis et al. 2015; Foster et al. 2016; Liu et al. 2017; Pacheco et al. 2018; dos Reis et al. 2018; Takezaki 2018), a practice that often generates controversy via widely differing time estimates (Battistuzzi et al. 2010; Christin et al. 2014; dos Reis et al. 2014; dos Reis et al. 2015; Foster et al. 2016; Liu et al. 2017; Pacheco et al. 2018; Takezaki 2018).

Therefore, we need a powerful method to accurately test whether evolutionary rates are autocorrelated in a phylogeny. Application of this method to molecular datasets representing taxonomic diversity across the tree of life will enable an assessment of the preponderance of autocorrelated rates in nature. Here, we introduce a new machine learning approach (CorrTest) that shows high power to detect autocorrelation between molecular rates. CorrTest is computationally efficient, and its application to a large number of datasets establishes the pervasiveness of rate autocorrelation in the tree of life.

## New Method

Machine learning (McL) is widely used to solve problems in many fields, including ecology (Christin et al. 2018; Willcock et al. 2018) and population genetics (Saminadin-Peter et al. 2012; Schrider and Kern 2016; Schrider and Kern 2018). We present a supervised machine learning (McL) framework (Bzdok et al. 2018) to build a predictive model that distinguishes between ABR and IBR models, which is a challenge in molecular phylogenetics and phylogenomics. In our McL approach, the input is a molecular phylogeny with branch lengths and the output is a classification that corresponds to whether or not the evolutionary rates in the phylogeny are autocorrelated among branches (ABR or IBR, respectively). An overview of our McL approach is presented in figure 1.

**Figure 1.**
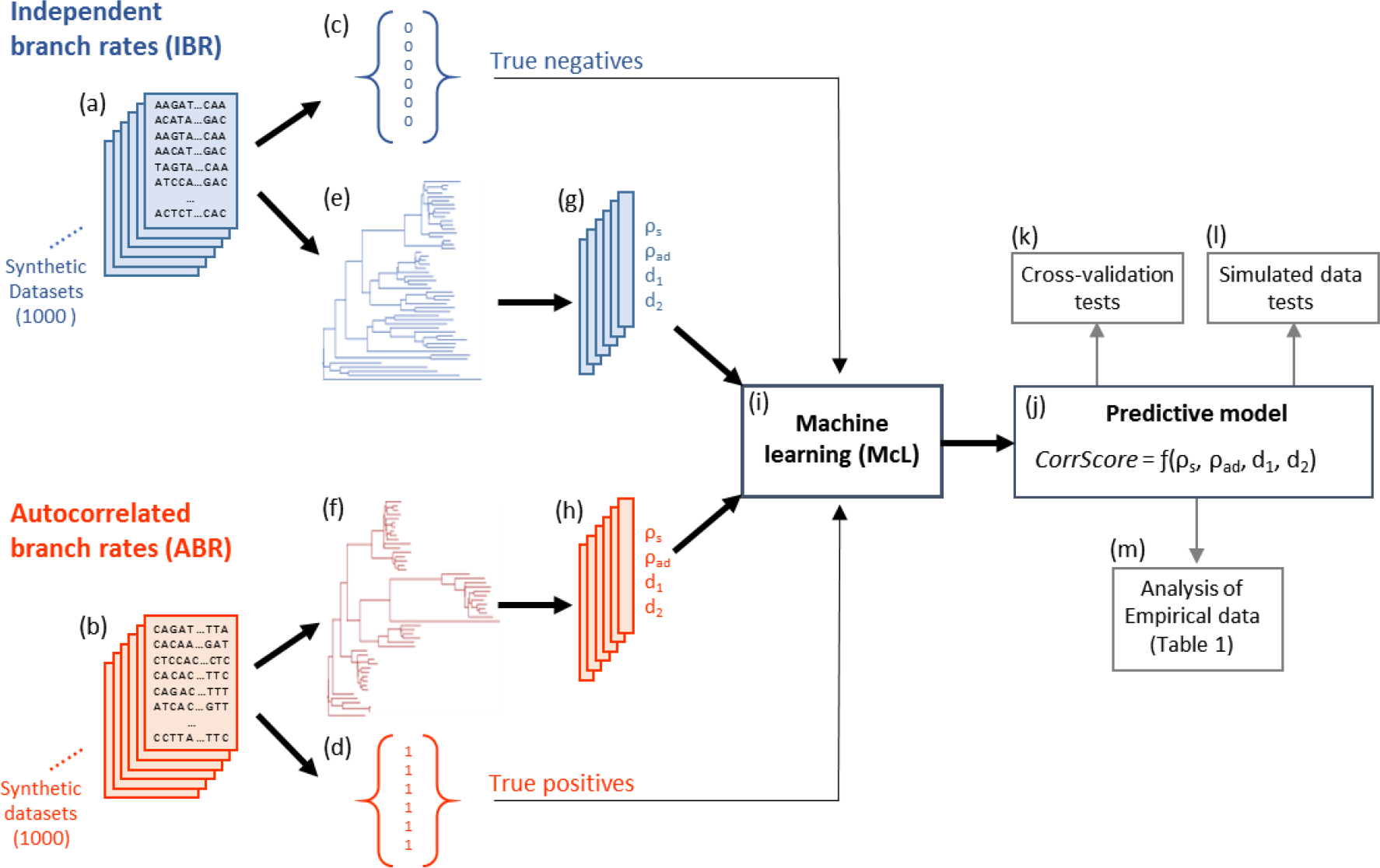
A flowchart showing an overview of the machine learning (McL) approach applied to develop the predictive model (CorrTest). We generated **(a)** 1,000 synthetic datasets that were evolved using an IBR model and **(b)** 1,000 synthetic datasets that were evolved using an ABR model. The numerical label (**c**) for all IBR datasets was 0 and (**d**) for all ABR datasets was 1. For each dataset, we estimated a molecular phylogeny with branch lengths (**e** and **f**) and computed ρ_s_, ρ_ad_, d_1_, and d_2_(**g** and **h**) that served as features during the supervised machine learning. **(i)** Supervised machine learning was used to develop a predictive relationship between the input features and labels. **(j)** The predictive model produces a CorrScore for an input phylogeny with branch lengths. The predictive model was **(k)** validated with 10-fold and 2-fold cross-validation tests, **(l)** tested using external simulated data, and then **(m)** applied to empirical data to examine the prevalence of rate autocorrelation in the tree of life.

To build a predictive model, McL needs measurable properties (features) that can be derived from the input data (phylogeny with branch lengths). The selection of informative and discriminating features (Fig. 1g and h) is critical for the success of McL. We derive relative lineage rates using a given molecular phylogeny with branch (“edge”) lengths (Fig. 1e and 1f) by using Tamura et al.’s method (Tamura et al. 2018), and use these lineage rates to generate informative features. This is necessary because we cannot derive branch rates without knowing node times in the phylogeny. For example, we need to know node times *t*_i_’s in figure 2 to convert branch lengths into branch rates, but these node times are what investigators wish to estimate by using a Bayesian approach which requires selection of a rate model. In contrast, estimation of relative lineage rates does not require the knowledge of divergence times, because an evolutionary lineage includes all the branches in the descendant subtree (e.g., lineage *a* contains branches with lengths *b*_1_, *b*_2_, and *b*_5_ in figure 2) and the relative rate between sister lineages is simply the ratio of the evolutionary depths of the two lineages (Tamura et al. 2018). In figure 2, *R*_a_ and *R*_b_ are two lineage rates, whose relative value can be estimated by the ratio of *L*_a_ and *L*_b_. Tamura et al. (2018) presented the relative rate framework (RRF) to estimate these relative lineage rates analytically by using branch lengths only. Furthermore, Tamura et al. (2018)’s method generates relative lineage rates such that all the lineage rates in a phylogeny are relative to the rate of the ingroup root lineage (*R*_0_, Fig. 2). This enables us to develop a number of features for building a McL predictive model.

**Figure 2.**
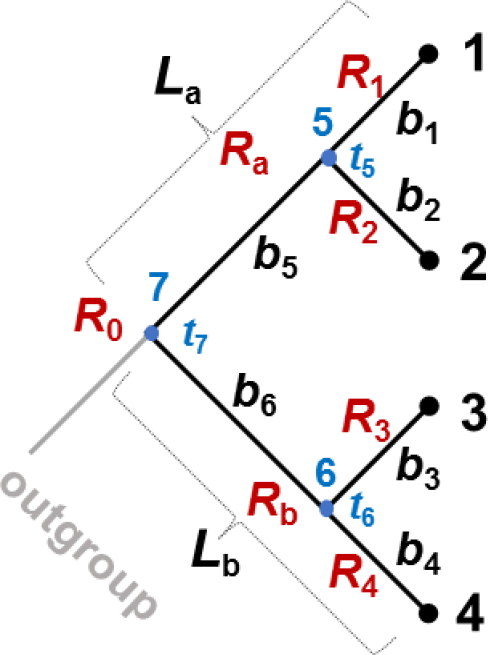
An evolutionary tree showing branch lengths (*b*), lineage lengths (*L*), lineage rates (*R*), and node times (*t*). Relative linage rates are computed from branch lengths using equations [28] - [31] and [34] - [39] in Tamura et al. (2018). Node times and branch rates are not required for estimating relative lineage rates.

We considered correlation between ancestral and descendant lineage rates (ρ_ad_), the correlation between the sister lineage rates (ρ_s_), and the decay in ρ_ad_ when one or two parents are skipped (d_1_ and d_2_, respectively) (see *Materials and Methods*). ρ_ad_ was considered as a feature because our analyses of simulated data showed that ρ_ad_ was much higher for phylogenetic trees in which molecular sequences evolved under an ABR model (0.96) than the IBR model (0.54, Fig. 3a). Importantly, ρ_ad_ is not expected to be zero under the IBR model because ρ_ad_ is a correlation between ancestral and descendant lineages, not branches, and the evolutionary rate of an ancestral lineage depends on the evolutionary rates of its descendant lineages (Tamura et al. 2018). While ρ_ad_ is greater than zero, it showed distinct patterns for ABR and IBR models and is, thus, a good candidate feature for McL (Fig. 3a). As our second feature, we considered the correlation between the sister lineages (ρ_s_), because ρ_s_ was higher for the ABR model (0.89) than the IBR model (0.00, Fig. 3b). Two additional features considered were the decay in ρ_ad_ when one or two parent branches are skipped (d_1_ and d_2_, respectively). We expect that ρ_ad_ will decay slower under ABR than IBR, which was confirmed (Fig. 3c). The selected set of candidate features (ρ_s_, ρ_ad_, d_1_, and d_2_) can be measured for any phylogeny with branch lengths, e.g., derived from molecular data using the Maximum Likelihood method. They are then used to train the machine learning classifier (Fig. 1i and j). For this purpose, we need a large set of phylogenies in which branch rates are autocorrelated (ABR = 1, Fig. 1d) and phylogenies in which the branch rates are independent (IBR = 0, Fig. 1c).

**Figure 3.**
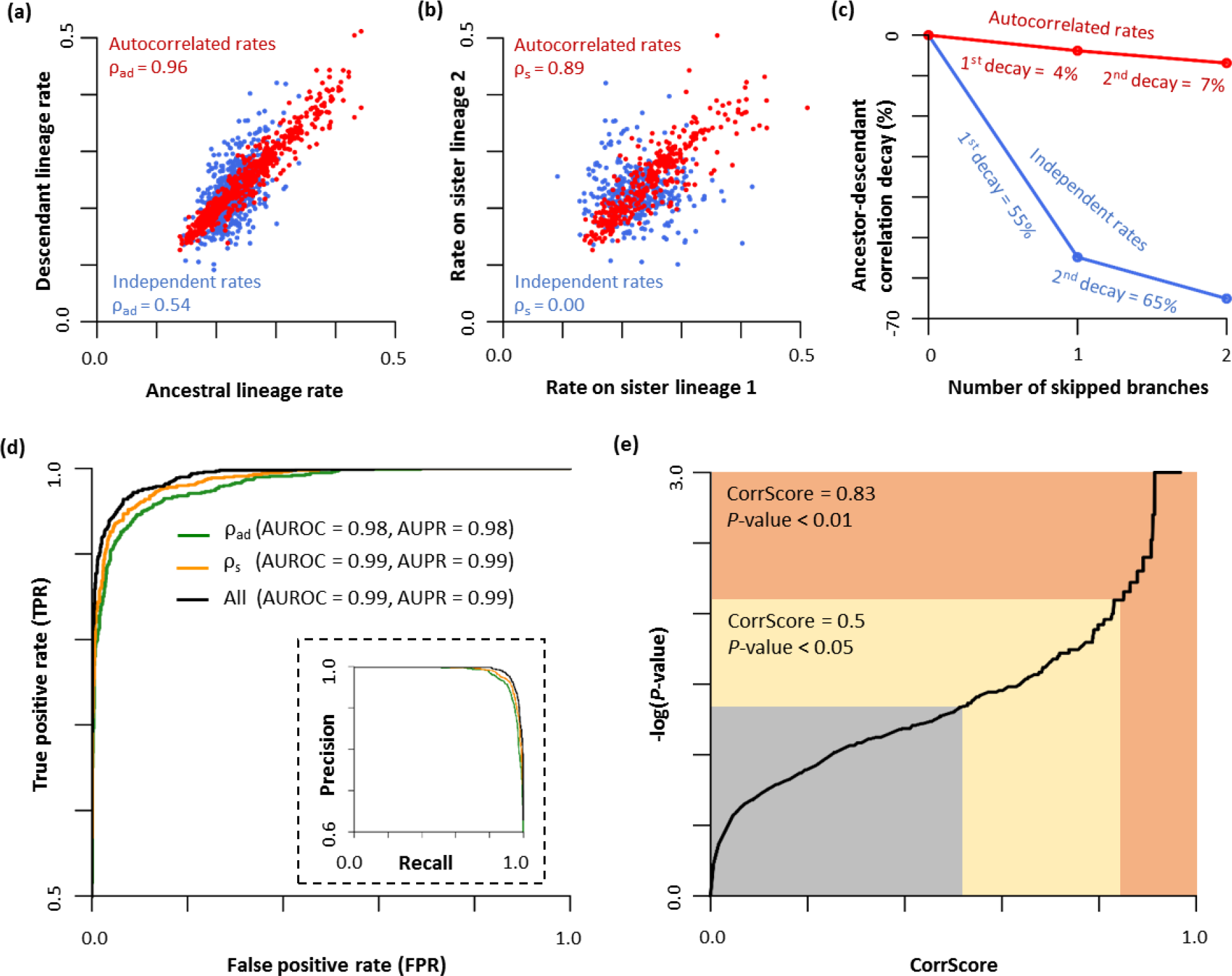
The relationship of **(a)** ancestral and direct descendant lineage rates and **(b)** sister lineage rates when the simulated evolutionary rates were autocorrelated with each other (red) or varied independently (blue). The correlation coefficients are shown. **(c)** The decay of correlation between ancestral and descendant lineages when we skip one intervening branch (1^st^ decay, d_1_) and when we skip two intervening branches (2^nd^ decay, d_2_). Percent decay values are shown. **(d)** Receiver Operator Characteristic (ROC) and Precision Recall (PR) curves (inset) of CorrTest for detecting branch rate model by using only ancestor-descendant lineage rates (ρ_ad_, green), only sister lineage rates (ρ_s_, orange), and all four features (all, black). The area under the curve is provided. **(e)** The relationship between the CorrScore produced by the machine learning model and the *P-*value. Independent rate model can be rejected when the CorrScore is greater than 0.83 at a significant level of *P* < 0.01, or when the CorrScore is greater than 0.5 at *P* < 0.05.

However, there is a paucity of empirical data for which ABR and IBR rates are firmly established. We, therefore, trained our McL model on a simulated dataset, a common practice in machine learning applications when reliable real world training datasets are few in number (Saminadin-Peter et al. 2012; Schrider and Kern 2016; Ekbatani et al. 2017; Le et al. 2017). We used computer simulations to generate 1,000 molecular datasets that evolved with ABR models and 1,000 molecular datasets that evolved with IBR models (Fig. 1a and b). To ensure the general utility of our model for analyses of diverse data, we simulated molecular sequences with varying numbers of species, degrees of rate autocorrelation, diversity of evolutionary rate and substitution pattern parameters (see *Materials and Methods*). Candidate features were computed for all 2,000 training datasets (Fig. 1g and h), each of which was associated with a numerical output state (IBR = 0 and ABR = 1; Fig. 1c and d). These features were used to build a predictive model by employing logistic regression (Fig. 1j). This predictive model was then used to generate a correlation score (CorrScore) for any phylogeny with branch lengths.

We also developed a conventional statistical test (CorrTest) based on CorrScore, which provides a *P*-value to decide whether the IBR model should be rejected. A high CorrScore indicates a high probability that the branch rates are autocorrelated. At a CorrScore greater than 0.5, Type I error (rejecting IBR when it was true) was less than 5%. Type I error of 1% (*P*-value of 0.01) was achieved with a CorrScore greater than 0.83 (Fig. 3e). CorrTest is available at Github (https://github.com/cathyqqtao/CorrTest) and in MEGA X (Kumar et al. 2018).

## RESULTS

We evaluated the sensitivity and specificity of our predictive model using receiver operating characteristic (ROC) curves, which measured the sensitivity of our method to detect rate autocorrelation when it is present (True Positive Rate, TPR) and when it was not present (False Positive Rate, FPR) at different CorrScore thresholds. The ROC curve for McL using all four features was the best, which led to the inclusion of all four features in the predictive model (Fig. 3d; *Material and Methods*). The area under the ROC (AUROC) was 99%, with a 95% TPR (i.e., ABR detection) achieved at the expense of only 5% FPR (Fig. 3d, black line). The area under the precision recall (AUPR) curve was also extremely high (0.99; Fig. 3d inset), which suggested that that CorrTest detects the presence of rate autocorrelation with very high accuracy and precision.

We also performed standard cross-validation tests using the simulated data to evaluate the accuracy of the predictive models when only a subset of data are used for training. In 10-fold cross-validation, the predictive model was developed using 90% of the synthetic datasets, and then its performance was tested on the remaining 10% of the datasets. The AUROC was greater than 0.99 and the accuracy was high (>94%). Even in the 2-fold cross-validation, where only half of the datasets (500 ABR and 500 IBR datasets) were used for training the model and the remaining half were used for testing, the AUROC was greater than 0.99 and the classification accuracy was greater than 92%. This suggested that the predictive model is robust.

We tested the performance of CorrTest on a large collection of simulated datasets where the correct rate model is known (Fig. 1l). In these datasets (Tamura et al. 2012), a different software and simulation schemes were used to generate sequences with a wide range of empirically derived G+C contents, transversion/transition ratios, and evolutionary rates under both ABR and IBR models (see *Materials and Methods*). CorrTest accuracy was greater than 94% in detecting ABR and IBR models correctly for datasets that were simulated with low and high G+C contents (Fig. 4a), small and large substitution rate biases (Fig. 4b), and different rates of evolution (Fig. 4c). As expected, CorrTest performed best on datasets that contain more and longer sequences (Fig. 4d).

**Figure 4.**
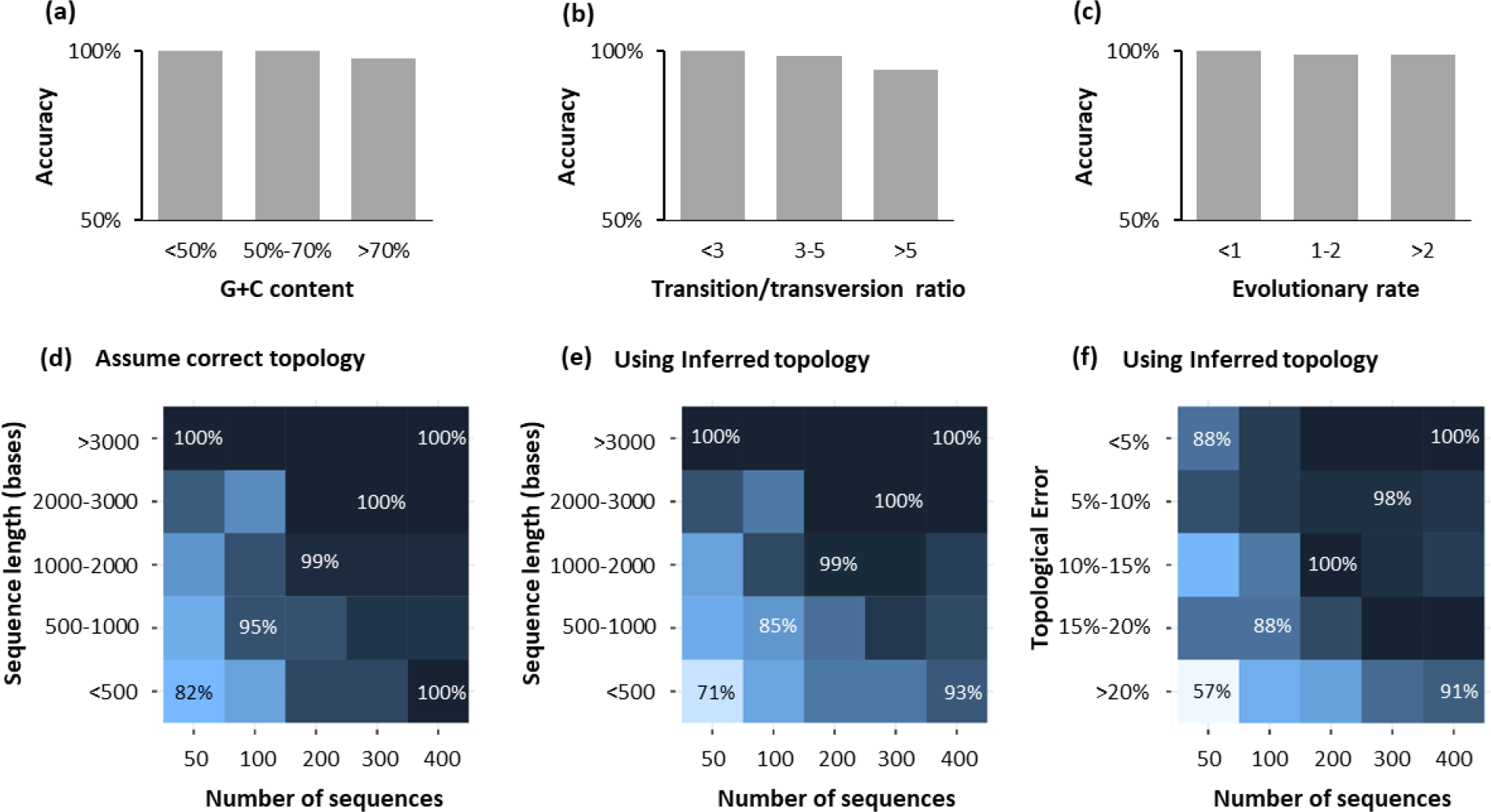
The performance of CorrTest in detecting autocorrelation and independent rate models in the analysis of datasets (Tamura et al. 2012) that were simulated with different **(a)** G+C contents, **(b)** transition/transversion rate ratios, and **(c)** average molecular evolutionary rates. Darker color indicates higher accuracy. The evolutionary rates are in the units of 10^−3^ substitutions per site per million years. (**d** - **f**) Patterns of CorrTest accuracy for datasets containing increasing number of sequences. The accuracy of CorrTest for different sequence length is shown when (**d**) the correct topology was assumed and (**e**) the topology was inferred. (**f**) The accuracy of CorrTest for datasets in which the inferred the topology contained small and large number of topological errors.

In the above analyses, we used the correct tree topology and nucleotide substitution model. We relaxed this requirement and evaluated CorrTest by first inferring a phylogeny using the Neighbor-joining method (Saitou and Nei 1987) with an oversimplified substitution model (Kimura 1980). Naturally, many inferred phylogenies contained topological errors, but we found the accuracy of CorrTest to be high as long as the dataset contained >100 sequences of length >1,000 base pairs (Fig. 4e). CorrTest also performed well when 20% of the tree partitions were incorrect in the inferred phylogeny (Fig. 4f). Therefore, CorrTest will be most reliable for large datasets, and is relatively robust to errors in phylogenetic inference.

### CorrTest versus Bayes factor analysis

We compared the performance of CorrTest with that of the Bayes factor approach. Because the Bayes factor method is computationally demanding, we limited our comparison to 100 datasets containing 100 sequences each (see *Material and Methods*). We computed Bayes factors (BF) by using the stepping-stone sampling (SS) method (see *Materials and Methods*). BF-SS analysis detected autocorrelation (*P* < 0.05) for 33% of the autocorrelated rate datasets (Fig. 5a, red curve in the ABR zone). This is because, for these datasets, the marginal log-likelihoods under the ABR model for 67% were very similar to or lower than that for the IBR model. Therefore, BF was conservative in rejecting the IBR model, as has been reported before (Ho et al. 2015). CorrTest performed better, it correctly detected the ABR model for 88% of the datasets (*P* < 0.05; Fig. 5b, red curve in ABR zone). For datasets that evolved with IBR model, BF-SS correctly detected the IBR model for 89% (Fig. 5a, blue curves in the IBR zone), whereas CorrTest correctly detected 86% (Fig. 5b, blue curve in the IBR zone). Therefore, BF performs well in correctly classifying phylogenies that evolved under IBR, but not ABR. The power of CorrTest to correctly infer ABR is responsible for its higher overall accuracy (87% vs. 61% for BF). Such a difference in accuracy was observed at different levels of statistical significance (Fig. 5c), for datasets that evolved with high and low rate autocorrelation (Fig. 5d), and for datasets that were simulated with low and high degrees of independent rate variation (Fig. 5e). These comparisons suggest that the machine learning method enables highly accurate detection of rate correlation in a given phylogeny and presents a computationally feasible alternative to Bayes Factor analyses for large datasets.

**Figure 5.**
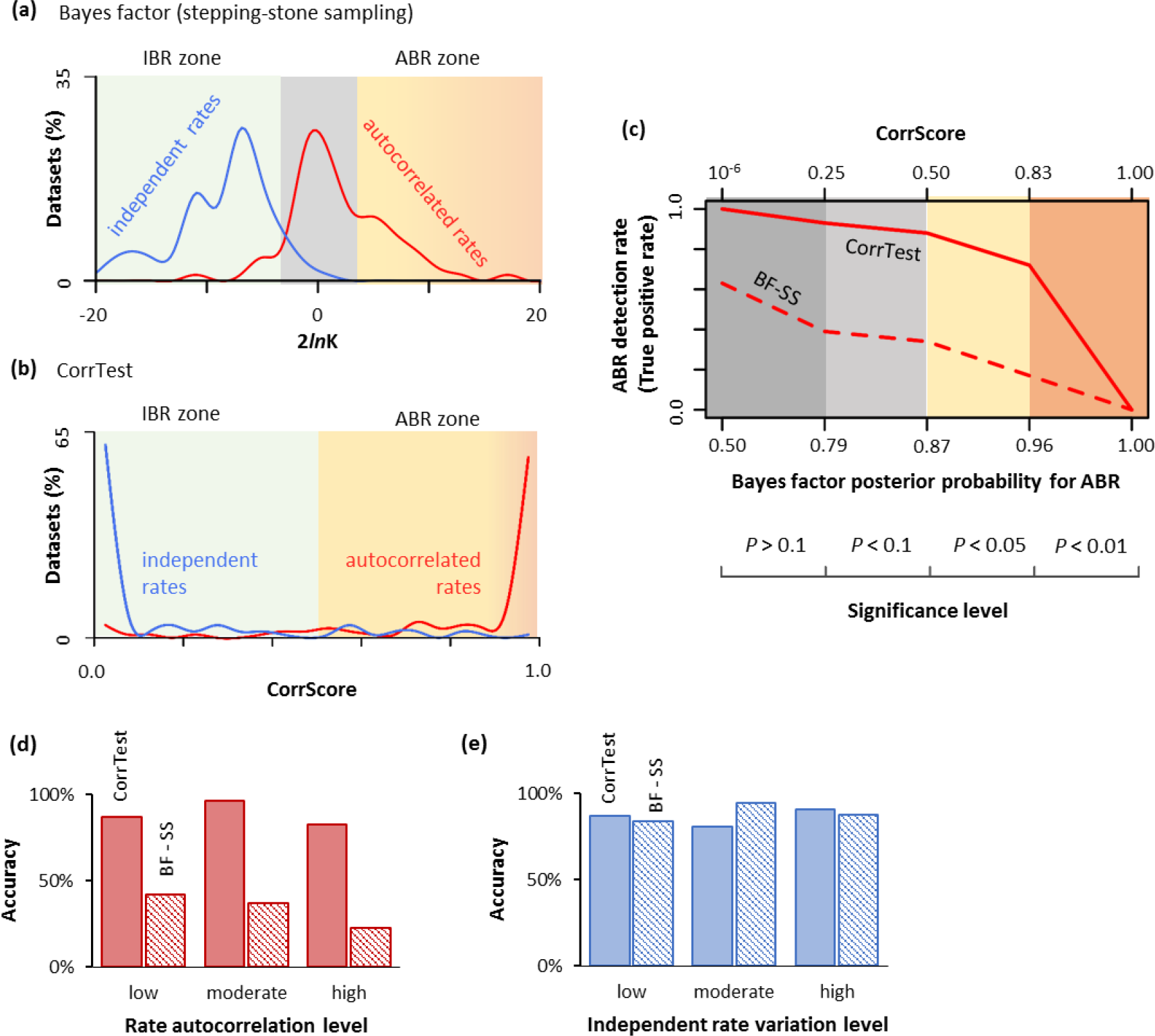
Comparisons of the performance of CorrTest and Bayes Factor analyses. **(a)** Distributions of 2 times the differences of marginal log-likelihood (2*ln*K) estimated via stepping-stone sampling method for datasets that were simulated with autocorrelated branch rates (ABR, red) and independent branch rates (IBR, blue). ABR is preferred (*P* < 0.05) when 2*ln*K is greater than 3.841 (ABR zone), and IBR is preferred when 2lnK is less than −3.841 (IBR zone). When 2*ln*K is between −3.841 and 3.841, the fit of the two rate models is not significantly different (gray shade). **(b)** The distributions of CorrScores in analyses of ABR (red) and IBR (blue) datasets. Rates are predicted to be autocorrelated if the CorrScore is greater than 0.5 (*P* < 0.05, ABR zone) and vary independently if the CorrScore is less than 0.5 (IBR zone). **(c)** The rate of detecting ABR model correctly (True Positive Rate) at different levels of statistical significance in Bayes factor (BF-SS) and CorrTest analyses. Posterior probabilities for ABR in BF-SS analysis are derived using the log-likelihood patterns in panel a. CorrTest *P*-values are derived using the CorrScore pattern in panel b. **(d)** The accuracy of identifying ABR model for datasets simulated with low (*v* < 0.1), moderate (0.1 ≤ *v* < 0.2), and high (*v* ≥ 0.2) levels of rate autocorrelation in Kishino et al. (2001)’s model. **(e)** The accuracy of identifying IBR model for datasets simulated at different degree of rate variation in Drummond et al. (2006): low (standard deviation < 0.2), moderate (0.2 ≤ standard deviation < 0.3), and high (standard deviation ≥ 0.3).

### Correlation of rates is common in molecular evolution

The high accuracy and fast computational speed of CorrTest enabled us to test the presence of autocorrelation in 16 large datasets from 12 published studies of eukaryotic species and 2 published studies of prokaryotic species encompassing diverse groups across the tree life. This included nuclear, mitochondrial and plastid DNA, and protein sequences from mammals, birds, insects, metazoans, plants, fungi, and prokaryotes (Table 1). CorrTest rejected the IBR model for all datasets (*P* < 0.05). In these analyses, we assumed a time-reversible process for base substitution. However, the violation of this assumption may produce biased results in phylogenetic analysis (Jayaswal et al. 2014). We, therefore, applied an unrestricted substitution model for analyzing all the nuclear datasets and confirmed that CorrTest rejected the IBR model in every case (*P* < 0.05). This robustness stems from the fact that the branch lengths estimated under the time-reversible and the unrestricted model are highly correlated for these data (*r*^2^ > 0.99). This is the reason why CorrTest produces reliable results even when an oversimplified model was used in analyzing the computer simulated data in which a complex model of sequence evolution was used to generate the sequence alignments (Fig. 4e and f).

**Table 1.** Patterns of rate autocorrelation inferred using the CorrTest approach.

| Taxonomic Group | Data type | Sequence count<sup>a</sup> | Sequence length | Substitution model | CorrTest score | P-value | 1/v<sup>b</sup> | Reference |
| --- | --- | --- | --- | --- | --- | --- | --- | --- |
| Mammals | Nuclear 4-fold degenerate sites | 138 | 1,671 | GTR + $\Gamma$ | 0.98 | < 0.001 | 3.21 | Meredith et al. (2011) |
| Mammals | Nuclear 3<sup>rd</sup> codon | 138 | 11,010 | GTR + $\Gamma$ | 0.99 | < 0.001 | 4.42 | Meredith et al. (2011) |
| Mammals | Nuclear proteins | 138 | 11,010 | JTT + $\Gamma$ | 0.99 | < 0.001 | 3.11 | Meredith et al. (2011) |
| Mammals | Mitochondrial DNA | 271 | 7,370 | HKY + $\Gamma$ | 0.98 | < 0.001 | 3.77 | dos Reis et al. (2012) |
| Birds | Nuclear DNA | 198 | 101,781 | GTR + $\Gamma$ | 1.00 | < 0.001 | 2.07 | Prum et al. (2015) |
| Birds | Nuclear 3<sup>rd</sup> codon positions | 222 | 1,364 | GTR + $\Gamma$ | 1.00 | < 0.001 | 2.11 | Claramunt et al. (2015) |
| Birds | Nuclear 1<sup>st</sup> and 2<sup>nd</sup> codon positions | 222 | 2,728 | GTR + $\Gamma$ | 1.00 | < 0.001 | 2.53 | Claramunt et al. (2015) |
| Insects | Nuclear proteins | 143 | 220,091 | LG + $\Gamma$ | 1.00 | < 0.001 | 8.68 | Misof et al. (2014) |
| Metazoans | Mitochondrial & nuclear proteins | 113 | 2,049 | LG + $\Gamma$ | 0.65 | < 0.05 | 40.0 | Erwin et al. (2011) |
| Plants | Plastid 3<sup>rd</sup> codon positions | 335 | 19,449 | GTR + $\Gamma$ | 1.00 | < 0.001 | 2.28 | Ruhfel et al. (2014) |
| Plants | Plastid proteins | 335 | 19,449 | JTT + $\Gamma$ | 1.00 | < 0.001 | 2.46 | Ruhfel et al. (2014) |
| Plants | Nuclear 1<sup>st</sup> and 2<sup>nd</sup> codon positions | 99 | 220,091 | GTR + $\Gamma$ | 1.00 | < 0.001 | 5.50 | Wickett et al. (2014) |
| Plants | Chloroplast and nuclear DNA | 124 | 5,992 | GTR + $\Gamma$ | 1.00 | < 0.001 | 2.64 | Beaulieu et al. (2015) |
| Fungi | Nuclear proteins | 85 | 609,772 | LG + $\Gamma$ | 0.97 | < 0.001 | 3.78 | Shen et al. (2016) |
| Prokaryotes | Nuclear proteins | 197 | 6,884 | JTT + $\Gamma$ | 0.79 | < 0.05 | 2.54 | Battistuzzi et al. (2009) |
| Prokaryotes | Nuclear proteins | 126 | 3,145 | JTT + $\Gamma$ | 0.83 | < 0.05 | 1.23 | Calteau et al. (2014) |

<sup>a</sup>Sequence count excludes the outgroup.
<sup>b</sup>1/v is the inverse of the autocorrelation parameter that is estimated by MCMCTree with the autocorrelated rate model in the time unit of 100My.

These results suggest that the correlation of rates among lineages is the rule, rather than the exception in molecular phylogenies. This pattern contrasts starkly with those reported in many previous studies (Drummond et al. 2006; Moore and Donoghue 2007; Brown et al. 2008; Bell et al. 2010; Smith et al. 2010; Linder et al. 2011; Jarvis et al. 2014; Lu et al. 2014; Barreda et al. 2015; Claramunt and Cracraft 2015; Prum et al. 2015; Feng et al. 2017; Barba-Montoya et al. 2018). In fact, all but three datasets (Battistuzzi and Hedges 2009; Erwin et al. 2011; Calteau et al. 2014) received very high prediction scores in CorrTest, resulting in extremely significant *P*-values (*P* < 0.001). The IBR model was also rejected for the other three datasets (*P* < 0.05), but their test scores were not as high, likely because they sparsely sample a large phylogenetic space. For example, the metazoan dataset (Erwin et al. 2011) contains sequences primarily from highly divergent species that shared common ancestors hundreds of millions of years ago. In this case, tip lineages in the phylogeny are long and their evolutionary rates are influenced by many un-sampled lineages. Such sampling effects weaken the rate correlation signal. We verified this behavior via an analysis of simulated data and found that CorrTest’s prediction scores decreased when taxon sampling and density were lowered (Fig. 6). Overall, CorrTest detected rate correlation in all the empirical datasets.

**Figure 6.**
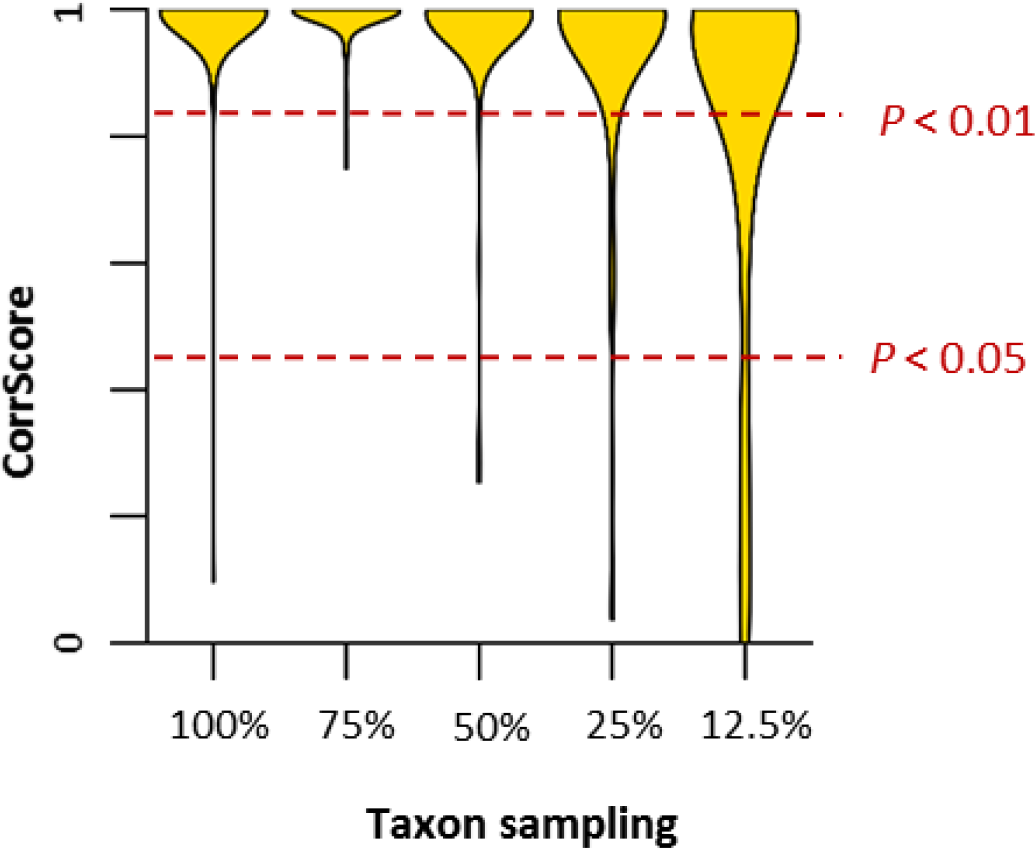
The distribution of CorrScore when data have different taxon sampling densities. The CorrScore decreases when the density of taxon sampling is lower, as there is much less information to discriminate between ABR and IBR. Red, dashed lines mark two statistical significance levels of 5% and 1%.

### Magnitude of the rate correlation in molecular data

CorrScore is influenced by the size of the dataset in addition to the degree of autocorrelation, so it is not a direct measure of the degree of rate autocorrelation (effect size) in a phylogeny. Instead, one should use a Bayesian approach to estimate the degree of rate correlation, for example, under Kishino et al. (2001)’s autocorrelated rate model. In this model, a single parameter (*v*) captures the degree of autocorrelation among branches in a phylogenetic tree. A low value of *v* indicates high autocorrelation, so, we use the inverse of *v* to represent the degree of rate autocorrelation. MCMCTree (Yang 2007) analyses of simulated datasets confirmed that the estimated *v* is linearly related to the true value (Fig. 7). In empirical data analyses, we find that the inverse of *v* is high for all datasets examined, which suggests ubiquitous high rate autocorrelation across the tree of life.

**Figure 7.**
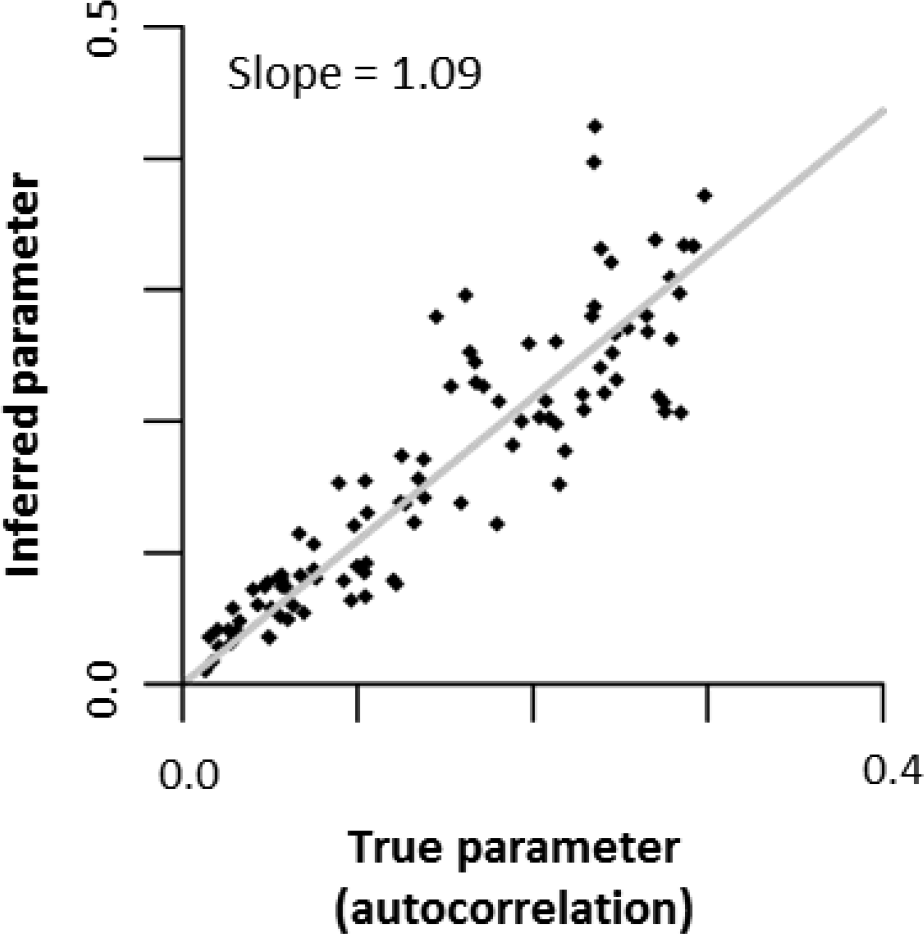
The relationship between the inferred autocorrelation parameter from MCMCTree and the true value. The gray line represents the best-fit regression line, which has a slope of 1.09.

Other interesting patterns emerge from this analysis. First, rate autocorrelation is highly significant not only for mutational rates (= substitution rate at neutral positions), which are expected to be similar in sister species because they inherit cellular machinery from a common ancestor, but also amino acid substitution rates, which are more strongly influenced by natural selection (Table 1). For example, synonymous substitution rates in the third codon positions and the four-fold degenerate sites in mammals (Meredith et al. 2011), which are largely neutral and are the best reflection of mutation rates (Kumar and Subramanian 2002), received high CorrScores of 0.99 and 0.98, respectively (*P* < 0.001). Second, our model also detected a strong signal of correlation for amino acid substitution rates in the same proteins (CorrScore = 0.99). Bayesian analyses showed that the degree of correlation is high in both cases: inverse of *v* was 3.21 in 4-fold degenerate sites and 3.11 in amino acid sequences. Third, mutational and substitution rates in both nuclear and mitochondrial genomes are highly autocorrelated (Table 1). These results establish that molecular and non-molecular evolutionary patterns are concordant, because morphological characteristics are also found to be similar between closely-related species (Sargis and Dagosto 2008; Lanfear et al. 2010; Cox and Hautier 2015) and correlated with taxonomic or geographic distance (Wyles et al. 1983; Shao et al. 2016).

## Discussion

Our results demonstrate that the machine learning framework is useful to develop a method to detect the presence of rate autocorrelation among branches in a phylogeny. This method yields CorrScore estimates that we found to be useful to develop a conventional statistical test, CorrTest, and generate an associated *P*-value. This model can be used for datasets with small and large number of sequences, because we already tested if higher CorrTest accuracy could be achieved by building specific predictive models that were trained separately by using data with ≤ 100 (M100), 100 – 200 (M200), 200 – 300 (M300), and > 300 (M400) sequences. A specific threshold was determined for each training subset and then was tested each model using Tamura et al. (2012)’s data with the corresponding number of sequences. For example, we used the threshold determined by the model trained with small data (≤ 100 sequences) on the test data that contain less than 100 sequences, and used the threshold determined by the model trained with large data (>300 sequences) on the large test data (400 sequences). We found that the accuracy of using the specific thresholds (Fig. 8) is similar to the accuracy when we used a global threshold (Fig. 4d - f). This is because the machine learning algorithm has automatically incorporated the impact of the number of sequences when it determined the relationship of four selected features (ρ_ad_, ρ_s_, d_1_ and d_2_). This justifies the usage of the globally trained CorrTest that we used in all the empirical analyses.

**Figure 8.**
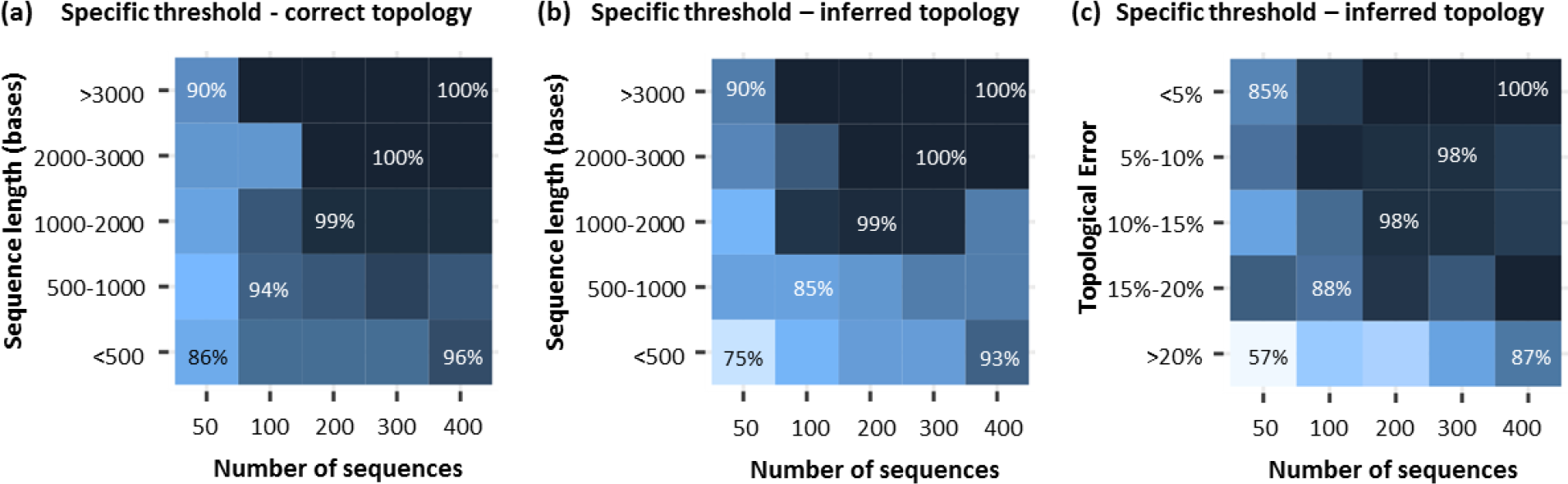
Patterns of CorrTest accuracy using M100, M200, M300, and M400 models for the corresponding test datasets (Tamura et al. 2012). Accuracies are shown for increasing number of sequences. The accuracy of CorrTest for different sequence length is shown when (**a**) the correct topology was assumed and (**b**) the topology was inferred. (**c**) The accuracy of CorrTest for datasets in which the inferred the topology contained small and large number of topological errors.

Our results suggest that the autocorrelated rate model should be the default in molecular clock analysis, and CorrTest can be used to test the independent rate model when sufficient numbers of sequences are available. Use of the autocorrelated rate model is important because model selection has a strong influence on the posterior credible intervals of divergence times (Battistuzzi et al. 2010). For example, the use of IBR model produces estimates of divergence time of two major groups of grasses that are 66% older (Christin et al. 2014) and origin of a major group of mammal (Erinaceidea) to be 30% older (Meredith et al. 2011) than estimates under ABR model. In fact, substantial differences between node age estimates under IBR and ABR models have been reported in many studies (Battistuzzi et al. 2010; Bell et al. 2010; Christin et al. 2014; dos Reis et al. 2015; Foster et al. 2016; Liu et al. 2017; Pacheco et al. 2018; Takezaki 2018). Thus, the use of an incorrect rate model has a large impact on time estimates, which may not always be alleviated by adding calibrations (Battistuzzi et al. 2010). Knowledge that evolutionary rates are generally autocorrelated within lineages will foster unbiased and precise dating of the tree of life, whenever one needs to choose a rate model to generate accurate Bayesian time estimates for use in studies of biodiversity, phylogeography, development, and genome evolution. However, it is important to appreciate that no single rate model may be adequate for Bayesian dating analyses, and one may need to use a mixture of models because different groups of species and genes in a large phylogeny may have evolved with different levels of autocorrelation (e.g., Lartillot et al. (2016)). In this sense, the results produced by CorrTest (and by Bayes Factor) analyses primarily detect the presence of rate autocorrelation, but they do not tell us if the rate autocorrelation exists in every clade of a phylogeny or if the degree of autocorrelation is the same in all the clades. One may apply CorrTest to individual clades (subtrees) to evaluate these patterns. For example, we applied CorrTest on subtrees to detect the existence of clade specific rate autocorrelation by dividing a few large empirical phylogenies (Meredith et al. 2011; dos Reis et al. 2012; Misof et al. 2014; Prum et al. 2015) into subtrees with at least 50 sequences. We found rate autocorrelation to be present in a vast majority of data subsets (results not shown), which supports the conclusion that rate autocorrelation is a common feature of molecular evolution of DNA and amino acid sequences.

## Materials and Methods

### Machine learning (McL) model

#### Training data for McL

We simulated nucleotide alignments using independent branch rate (IBR) and autocorrelated branch rate (ABR) models using the NELSI package (Ho et al. 2015) with a variety of empirically drived parameters or parameters that were used in a previous study. In IBR, branch-specific rates were drawn from a lognormal distribution with a mean gene rate and a standard deviation (in log-scale) that varied from 0.1 to 0.4, previously used in a study simulating independent rates with different levels of variation (Ho et al. 2015). In ABR, branch-specific rates were simulated under an autocorrelated process (Kishino et al. 2001) with an initial rate set as the mean rate derived from an empirical gene and an autocorrelated parameter, *v*, that was randomly chosen from 0.01 to 0.3, previously used in a study simulating low, moderate and high degrees of autocorrelated rates (Ho et al. 2015). We used SeqGen (Grassly et al. 1997) to generate alignments under Hasegawa-Kishino-Yano (HKY) model (Hasegawa et al. 1985) with 4 discrete gamma categories by using a master phylogeny, consisting of 60-400 ingroup taxa randomly sampled from the bony-vertebrate clade in the Timetree of Life (Hedges and Kumar 2009). Mean evolutionary rates, G+C contents, transition/transversion ratios and numbers of sites for simulation were derived from empirical distributions (Rosenberg and Kumar 2003). 1,000 molecular datasets were generated under ABR and IBR model separately and these 2,000 simulated datasets were used as training data in building the machine learning model.

#### Calculation of features for McL

Lineage-specific rate estimates (*R*_i_’s) were obtained using equations [28] - [31] and [34] - [39] in Tamura et al. (2018). For any given node in the phylogeny, we extracted the relative rates of its ancestral clade (*R*_a_) and two direct descendant clades (*R*_1_ and *R*_2_). Then, we calculated correlation between ancestral lineage and its direct descendant lineage rate to obtain estimates of ancestor-descendant rate correlation (ρ_ad_). We also calculated correlation between sister lineage rates (ρ_s_), for which the lineage rates of sister pairs are randomly labeled. The labeling of sister pairs have small impact on ρ_s_ when the number of sequences in the phylogeny is not too small (>50). For smaller datasets, we found that it is best to generate multiple ρ_s_ estimates, each using randomly labelled sister pairs, in order to eliminate any bias that may result from the arbitrary designation of sister pairs. In this case, we use the mean *ρ*_s_ from multiple replicates in the CorrTest analysis. To avoid the assumption of linear correlation between lineages, we used Spearman rank correlation because it can capture both linear and non-linear correlation between two vectors. Two additional features included in McL measure the decay in ρ_ad_ when one or two intervening branches are skipped (d_1_ and d_2_, respectively). We first estimated ρ_ad_skip1_as the correlation between rates where the ancestor and descendant were separated by one intervening branch, and ρ_ad_skip2_ as the correlation between rates where the ancestor and descendant were separated by two intervening branches. This skipping reduces ancestor-descendant correlation, which we then used to derive the decay of correlation values by using equations d_1_ = (ρ_ad_ - ρ_ad_skip1_)/ρ_ad_ and d_2_ = (ρ_ad_ − ρ_ad_skip2_)/ρ_ad_. These two features improved the accuracy of our model slightly. In the analyses of empirical datasets, we found that a large amount of missing data (>50%) can result in unreliable estimates of branch lengths and other phylogenetic errors (Wiens and Moen 2008; Lemmon et al. 2009; Filipski et al. 2014; Xi et al. 2015; Marin and Hedges 2018). In this case, we recommend computing selected features using only those lineage pairs for which >50% of the positions contain valid data, or remove sequences with a large amount of missing data.

#### Building the McL predictive model

We trained a logistic regression model using the skit-learn module (Pedregosa et al. 2011), which is a python toolbox for data mining and data analysis using machine learning algorithms, with only ρ_ad_, only ρ_s_ or all 4 features (ρ_s_, ρ_ad_, d_1_ and d_2_) using 2,000 simulated training datasets (1,000 with ABR model and 1,000 with IBR model). For each training data, we inferred the branch lengths from the molecular sequences with a fixed topology first and used these inferred branch lengths to estimate relative lineage rates for computing all the selected features. A response value of 1 was given to true positive cases (autocorrelated rates) and 0 was assigned to true negative cases (independent rates). Thus, the prediction scores (CorrScore) were between 0 and 1. A high score representing a higher probability that the rates are autocorrelated. Then the thresholds at 5% and 1% significant levels can be determined.

### Test datasets

Tamura et al. (2012)’s simulated data were used to evaluate CorrTest’s performance. We present the test results for the data simulated using ABR model (autocorrelated lognormal distribution) and IBR model (independent uniform distribution with 50% rate variation) here. We tested the performance of our model on ABR and IBR data with different GC contents, transition/transversion ratios, and evolutionary rates. We randomly sampled 50, 100, 200, and 300 sequences from the original 400 sequences and conducted CorrTest using the correct, error-prone topology inferred by the Neighbor-joining method (Saitou and Nei 1987) with an oversimplified substitution model (Kimura 1980). We also tested CorrTest’s performance on data simulated under an IBR model process with 100% rate variation and found that CorrTest works perfectly (100% accuracy; results not shown). In addition, we conducted another set of simulations using IBR (independent lognormal distribution) and ABR (autocorrelated lognormal distribution) (Kishino et al. 2001) model with 100 replicates each using the same strategy as a training data simulation (described above) on a master phylogeny of 100 taxa randomly sampled from the bony-vertebrate clade in the Timetree of Life (Hedges and Kumar 2009). These 200 datasets were used to conduct CorrTest and Bayes factor analyses and to obtain the autocorrelation parameter (*v*) in MCMCTree (Yang 2007).

### CorrTest analyses

All CorrTest analyses were conducted using a customized R code (available from https://github.com/cathyqqtao/CorrTest). We estimated branch lengths of a tree topology on sequence alignments using maximum likelihood method (or Neighbor-Joining method when we tested the robustness of our model to topological error) in MEGA (Kumar et al. 2012; Kumar et al. 2016). Then we used those estimated branch lengths to compute relative lineages rates using RRF (Tamura et al. 2012; Tamura et al. 2018) and calculated the value of selected features (*ρ*_s_, *ρ*_ad_, d_1_ and d_2_) to obtain the CorrScore. We conducted CorrTest on the CorrScore to estimate the *P*-value of detecting rate autocorrealtion. No calibration was needed for CorrTest analyses.

### Bayes factor analyses

We computed the Bayes factor via stepping-stone sampling (BF-SS) (Xie et al. 2011) with n = 20 and a = 5 using mcmc3r package (dos Reis et al. 2018). We chose BF-SS because the harmonic mean estimator has many statistical shortcomings (Lepage et al. 2007; Xie et al. 2011; Baele et al. 2013) and thermodynamic integration (Silvestro et al. 2011; dos Reis et al. 2018) is less efficient than BF-SS. For each dataset, we computed the log-likelihoods (*ln*K) of using IBR model and ABR model. The Bayes factor posterior probability for ABR was calculated as shown in dos Reis et al. (2018). We used only one calibration point at the root (true age with a narrow uniform distribution) in all the Bayesian analyses, as it is the minimum number of calibrations required by MCMCTree (Yang 2007). For other priors, we used diffused distributions of “rgene_gamma = 1 1”, “sigma2_gamma=1 1” and “BDparas = 1 1 0”. In all Bayesian analyses, two independent runs of 5,000,000 generations each were conducted, and results were checked in Tracer (Rambaut et al. 2014) for convergence. ESS values were higher than 200 after removing 10% burn-in samples for each run.

### Analysis of empirical datasets

We used 16 datasets from 12 published studies of eukaryotes and 2 published studies of prokaryotes that cover the major groups in the tree of life (Table 1). These were selected because they did not contain too much missing data (<50%) and represented >80 sequences. As we know, a large amount of missing data (>50%) can result in unreliable estimates of branch lengths and other phylogenetic errors (Wiens and Moen 2008; Lemmon et al. 2009; Filipski et al. 2014; Xi et al. 2015; Marin and Hedges 2018) and potentially bias CorrTest result. When a phylogeny and branch lengths were available from the original study, we estimated relative rates directly from the branch lengths via the relative rate framework (Tamura et al. 2018) and computed selected features to conduct CorrTest. Otherwise, maximum likelihood estimates of branch lengths were obtained in MEGA (Kumar et al. 2012; Kumar et al. 2016) using the published topology, sequence alignments, and the substitution model specified in the original article.

To obtain the autocorrelation parameter (*v*), we used MCMCTree (Yang 2007) with the same input priors as the original study, but no calibration priors were used in order to avoid the undue influence of calibration uncertainty densities on the estimate of *v*. We did, however, provide a root calibration because MCMCTree requires it. For this purpose, we used the root calibration provided in the original article or selected the median age of the root node in the TimeTree database (Hedges et al. 2006; Kumar et al. 2017) ± 50My (soft uniform distribution) as the root calibration, as this does not impact the estimation of *v*. Bayesian analyses required long computational times, so we used the original alignments in MCMCTree to infer *v* if alignments were shorter than 20,000 sites. If the alignments were longer than 20,000 sites, we randomly selected 20,000 sites from the original alignments. However, one dataset (Ruhfel et al. 2014) contained more than 300 ingroup species, such that even alignments of 20,000 sites required prohibitive amounts of memory. In this case, we randomly selected 2,000 sites from the original alignments to use in MCMCtree for *v* inference (similar results were obtained with a different site subset). Two independent runs of 5,000,000 generations each were conducted, and results were checked in Tracer (Rambaut et al. 2014) for convergence. ESS values were higher than 200 after removing 10% burn-in samples for each run. All empirical datasets are available at https://github.com/cathyqqtao/CorrTest.

## Acknowledgements

We thank Xi Hang Cao for assisting on building the machine learning model, and Drs. Bui Quang Minh, Beatriz Mello, Heather Rowe, Ananias Escalante, Maria Pacheco, and S. Blair Hedges for critical comments and editorial suggestions. This research was supported by grants from National Aeronautics and Space Administration (NASA NNX16AJ30G), National Institutes of Health (GM0126567-01; LM012487-03), National Science Foundation (NSF 1661218), and Tokyo Metropolitan University (DB105).

## References

Baele G, Li WLS, Drummond AJ, Suchard MA, Lemey P. 2013. Accurate model selection of relaxed molecular clocks in bayesian phylogenetics. Mol. Biol. Evol. 30:239–243.

Barba-Montoya J, Dos Reis M, Schneider H, Donoghue PCJ, Yang Z. 2018. Constraining uncertainty in the timescale of angiosperm evolution and the veracity of a Cretaceous Terrestrial Revolution. New Phytol. 218:819–834.

Barreda VD, Palazzesi L, Tellería MC, Olivero EB, Raine JI, Forest F. 2015. Early evolution of the angiosperm clade Asteraceae in the Cretaceous of Antarctica. Proc. Natl. Acad. Sci. U.S.A. 112:10989–10994.

Battistuzzi FU, Filipski A, Hedges SB, Kumar S. 2010. Performance of relaxed-clock methods in estimating evolutionary divergence times and their credibility intervals. Mol. Biol. Evol. 27:1289–1300.

Battistuzzi FU, Hedges SB. 2009. A major clade of prokaryotes with ancient adaptations to life on land. Mol. Biol. Evol. 26:335–343.

Beaulieu JM, O’Meara BC, Crane P, Donoghue MJ. 2015. Heterogeneous rates of molecular evolution and diversification could explain the Triassic age estimate for angiosperms. Syst. Biol. 64:869–878.

Bell CD, Soltis DE, Soltis PS. 2010. The age and diversification of the angiosperms re-revisited. Am. J. Bot. 97:1296–1303.

Brown JW, Rest JS, García-Moreno J, Sorenson MD, Mindell DP. 2008. Strong mitochondrial DNA support for a Cretaceous origin of modern avian lineages. BMC Biol. 6:6.

Buck CB, Van Doorslaer K, Peretti A, Geoghegan EM, Tisza MJ, An P, Katz JP, Pipas JM, McBride AA, Camus AC, et al. 2016. The Ancient Evolutionary History of Polyomaviruses. PLoS Pathog. 12:e1005574.

Bzdok D, Krzywinski M, Altman N. 2018. Machine learning: supervised methods. Nat. Methods 15:5–6.

Calteau A, Fewer DP, Latifi A, Coursin T, Laurent T, Jokela J, Kerfeld CA, Sivonen K, Piel J, Gugger M. 2014. Phylum-wide comparative genomics unravel the diversity of secondary metabolism in Cyanobacteria. BMC Genomics 15:977.

Christin P-A, Spriggs E, Osborne CP, Strӧmberg CA, Salamin N, Edwards EJ. 2014. Molecular dating, evolutionary rates, and the age of the grasses. Syst. Biol. 63:153–165.

Christin S, Hervet E, Lecomte N. 2018. Applications for deep learning in ecology. bioRxiv:334854.

Claramunt S, Cracraft J. 2015. A new time tree reveals Earth history’s imprint on the evolution of modern birds. Sci. Adv. 1:e1501005.

Cox PG, Hautier L. 2015. Evolution of the Rodents: Volume 5: Advances in Phylogeny, Functional Morphology and Development. (Cox, P.G. and Hautier, L., editor.). Cambridge: Cambridge University Press

Dos Reis M, Donoghue PC, Yang Z. 2016. Bayesian molecular clock dating of species divergences in the genomics era. Nat. Rev. Genet. 17:71–80.

Dos Reis M, Gunnell GF, Barba-Montoya J, Wilkins A, Yang Z, Yoder AD. 2018. Using phylogenomic data to explore the effects of relaxed clocks and calibration strategies on divergence time estimation: primates as a test case. Syst. Biol. 67:594–615.

Dos Reis M, Inoue J, Hasegawa M, Asher RJ, Donoghue PC, Yang Z. 2012. Phylogenomic datasets provide both precision and accuracy in estimating the timescale of placental mammal phylogeny. Proc. R. Soc. B 279:3491–3500.

Dos Reis M, Thawornwattana Y, Angelis K, Telford MJ, Donoghue PC, Yang Z. 2015. Uncertainty in the Timing of Origin of Animals and the Limits of Precision in Molecular Timescales. Curr. Biol. 25:1–12.

Dos Reis M, Zhu T, Yang Z. 2014. The impact of the rate prior on Bayesian estimation of divergence times with multiple loci. Syst. Biol. 64:555–565.

Drummond AJ, Ho SYW, Phillips MJ, Rambaut A. 2006. Relaxed phylogenetics and dating with confidence. PLoS Biol. 4:88–99.

Ekbatani HK, Pujol O, Segui S. 2017. Synthetic Data Generation for Deep Learning in Counting Pedestrians. In: Pattern Recognition Applications and Methods (ICPRAM), 2017 The International Conference on. p. 318–323.

Erwin DH, Laflamme M, Tweedt SM, Sperling EA, Pisani D, Peterson KJ. 2011. The Cambrian conundrum: early divergence and later ecological success in the early history of animals. Science 334:1091–1097.

Feng Y-J, Blackburn DC, Liang D, Hillis DM, Wake DB, Cannatella DC, Zhang P. 2017. Phylogenomics reveals rapid, simultaneous diversification of three major clades of Gondwanan frogs at the Cretaceous-Paleogene boundary. Proc. Natl. Acad. Sci. U.S.A. 114:E5864–E5870.

Filipski A, Murillo O, Freydenzon A, Tamura K, Kumar S. 2014. Prospects for building large timetrees using molecular data with incomplete gene coverage among species. Mol. Biol. Evol. 31:2542–2550.

Foster CS, Sauquet H, Van der Merwe M, McPherson H, Rossetto M, Ho SY. 2016. Evaluating the impact of genomic data and priors on Bayesian estimates of the angiosperm evolutionary timescale. Syst. Biol. 66:338–351.

Gillespie JH. 1984. The molecular clock may be an episodic clock. Proc. Natl. Acad. Sci. U.S.A. 81:8009–8013.

Grassly NC, Adachi J, Rambaut A. 1997. Seq-Gen: an application for the Monte Carlo simulation of protein sequence evolution along phylogenetic trees. Comput. Appl. Biosci. 13:235–238.

Hasegawa M, Kishino H, Yano T. 1985. Dating of the human-ape splitting by a molecular clock of mitochondrial DNA. J. Mol. Evol. 22:160–174.

Hedges SB, Dudley J, Kumar S. 2006. TimeTree: a public knowledge-base of divergence times among organisms. Bioinformatics 22:2971–2972.

Hedges SB, Kumar S. 2009. The Timetree of Life. New York: Oxford University Press.

Hertweck KL, Kinney MS, Stuart SA, Maurin O, Mathews S, Chase MW, Gandolfo MA, Pires JC. 2015. Phylogenetics, divergence times and diversification from three genomic partitions in monocots. Bot. J. Linn. Soc. 178:375–393.

Ho SY, Duchêne S, Duchêne D. 2015. Simulating and detecting autocorrelation of molecular evolutionary rates among lineages. Mol. Ecol. Resour. 15:688–696.

Ho SY, Duchêne S. 2014. Molecular-clock methods for estimating evolutionary rates and timescales. Mol. Ecol. 23:5947–5965.

Jarvis ED, Mirarab S, Aberer AJ, Li B, Houde P, Li C, Ho SY, Faircloth BC, Nabholz B, Howard JT, et al. 2014. Whole-genome analyses resolve early branches in the tree of life of modern birds. Science 346:1320–1331.

Jayaswal V, Wong TK, Robinson J, Poladian L, Jermiin LS. 2014. Mixture models of nucleotide sequence evolution that account for heterogeneity in the substitution process across sites and across lineages. Syst. Biol. 63:726–742.

Kimura M. 1980. A simple method for estimating evolutionary rates of base substitutions through comparative studies of nucleotide sequences. J. Mol. Evol. 16:111–120.

Kimura M. 1983. The neutral theory of molecular evolution. Cambridge: Cambridge University Press

Kishino H, Thorne JL, Bruno WJ. 2001. Performance of a divergence time estimation method under a probabilistic model of rate evolution. Mol. Biol. Evol. 18:352–361.

Kumar S, Hedges SB. 2016. Advances in time estimation methods for molecular data. Mol. Biol. Evol. 33:863–869.

Kumar S, Stecher G, Li M, Knyaz C, Tamura K. 2018. MEGA X: Molecular Evolutionary Genetics Analysis across Computing Platforms. Mol. Biol. Evol. 35:1547–1549.

Kumar S, Stecher G, Peterson D, Tamura K. 2012. MEGA-CC: computing core of molecular evolutionary genetics analysis program for automated and iterative data analysis. Bioinformatics 28:2685–2686.

Kumar S, Stecher G, Suleski M, Hedges SB. 2017. TimeTree: A Resource for Timelines, Timetrees, and Divergence Times. Mol. Biol. Evol. 34:1812–1819.

Kumar S, Stecher G, Tamura K. 2016. MEGA7: Molecular Evolutionary Genetics Analysis version 7.0 for bigger datasets. Mol. Biol. Evol. 33:1870–1874.

Kumar S, Subramanian S. 2002. Mutation rates in mammalian genomes. Proc. Natl. Acad. Sci. U.S.A. 99:803–808.

Kumar S. 2005. Molecular clocks: four decades of evolution. Nat. Rev. Genet. 6:654–662.

Lanfear R, Welch JJ, Bromham L. 2010. Watching the clock: studying variation in rates of molecular evolution between species. Trends Ecol. Evol. 25:495–503.

Lartillot N, Phillips MJ, Ronquist F. 2016. A mixed relaxed clock model. Phil. Trans. R. Soc. B 371:20150132.

Le TA, Baydin AG, Zinkov R, Wood F. 2017. Using synthetic data to train neural networks is model-based reasoning. In: Neural Networks (IJCNN), 2017 International Joint Conference on. p. 3514–3521.

Lemmon AR, Brown JM, Stanger-Hall K, Lemmon EM. 2009. The effect of ambiguous data on phylogenetic estimates obtained by maximum likelihood and Bayesian inference. Syst. Biol. 58:130–145.

Lepage T, Bryant D, Philippe H, Lartillot N. 2007. A general comparison of relaxed molecular clock models. Mol. Biol. Evol. 24:2669–2680.

Linder M, Britton T, Sennblad B. 2011. Evaluation of Bayesian models of substitution rate evolution-parental guidance versus mutual independence. Syst. Biol. 60:329–342.

Liu L, Zhang J, Rheindt FE, Lei F, Qu Y, Wang Y, Zhang Y, Sullivan C, Nie W, Wang J, et al. 2017. Genomic evidence reveals a radiation of placental mammals uninterrupted by the KPg boundary. Proc. Natl. Acad. Sci. U.S.A. 114:E7282–E7290.

Lu Y, Ran J-H, Guo D-M, Yang Z-Y, Wang X-Q. 2014. Phylogeny and divergence times of gymnosperms inferred from single-copy nuclear genes. PLoS One 9:e107679.

Lynch M. 2010. Evolution of the mutation rate. Trends Genet. 26:345–352.

Magallón S, Hilu KW, Quandt D. 2013. Land plant evolutionary timeline: gene effects are secondary to fossil constraints in relaxed clock estimation of age and substitution rates. Am. J. Bot. 100:556–573.

Marin J, Hedges SB. 2018. Undersampling genomes has biased time and rate estimates throughout the tree of life. Mol. Biol. Evol. 35:2077–2084.

Meredith RW, Janečka JE, Gatesy J, Ryder OA, Fisher CA, Teeling EC, Goodbla A, Eizirik E, Simão TL, Stadler T, et al. 2011. Impacts of the Cretaceous Terrestrial Revolution and KPg extinction on mammal diversification. Science 334:521–524.

Metsky HC, Matranga CB, Wohl S, Schaffner SF, Freije CA, Winnicki SM, West K, Qu J, Baniecki ML, Gladden-Young A, et al. 2017. Zika virus evolution and spread in the Americas. Nature 546:411–415.

Misof B, Liu S, Meusemann K, Peters RS, Donath A, Mayer C, Frandsen PB, Ware J, Flouri T, Beutel RG, et al. 2014. Phylogenomics resolves the timing and pattern of insect evolution. Science 346:763–767.

Moore BR, Donoghue MJ. 2007. Correlates of diversification in the plant clade Dipsacales: geographic movement and evolutionary innovations. Am. Nat. 170 Suppl 2:S28–55.

Pacheco MA, Matta NE, Valkiunas G, Parker PG, Mello B, Stanley CE, Lentino M, Garcia-Amado MA, Cranfield M, Kosakovsky Pond SL, et al. 2018. Mode and Rate of Evolution of Haemosporidian Mitochondrial Genomes: Timing the Radiation of Avian Parasites. Mol. Biol. Evol. 35:383–403.

Pedregosa F, Varoquaux G, Gramfort A, Michel V, Thirion B, Grisel O, Blondel M, Prettenhofer P, Weiss R, Dubourg V, et al. 2011. Scikit-learn: Machine learning in Python. J. Mach. Learn. Res. 12:2825–2830.

Prum RO, Berv JS, Dornburg A, Field DJ, Townsend JP, Lemmon EM, Lemmon AR. 2015. A comprehensive phylogeny of birds (Aves) using targeted next-generation DNA sequencing. Nature 526:569–578.

Rambaut A, Suchard M, Xie D, Drummond A. 2014. Tracer v1.6. Available from: http://beast.bio.ed.ac.uk/Tracer

Rosenberg MS, Kumar S. 2003. Heterogeneity of nucleotide frequencies among evolutionary lineages and phylogenetic inference. Mol. Biol. Evol. 20:610–621.

Ruhfel BR, Gitzendanner MA, Soltis PS, Soltis DE, Burleigh JG. 2014. From algae to angiosperms-inferring the phylogeny of green plants (Viridiplantae) from 360 plastid genomes. BMC Evol. Biol. 14:23.

Saitou N, Nei M. 1987. The neighbor-joining method: a new method for reconstructing phylogenetic trees. Mol. Biol. Evol. 4:406–425.

Saminadin-Peter SS, Kemkemer C, Pavlidis P, Parsch J. 2012. Selective sweep of a cis-regulatory sequence in a non-African population of Drosophila melanogaster. Mol. Biol. Evol. 29:1167–1174.

Sanderson MJ. 1997. A nonparametric approach to estimating divergence times in the absence of rate constancy. Mol. Biol. Evol. 14:1218–1231.

Sargis EJ, Dagosto M. 2008. Mammalian evolutionary morphology: a tribute to Frederick S. Szalay. (Sargis, Eric J. and Dagosto, Marian, editor.). Springer Netherlands

Schrider DR, Kern AD. 2016. S/HIC: Robust Identification of Soft and Hard Sweeps Using Machine Learning. PLoS Genet. 12:e1005928.

Schrider DR, Kern AD. 2018. Supervised Machine Learning for Population Genetics: A New Paradigm. Trends Genet. 34:301–312.

Shao S, Quan Q, Cai T, Song G, Qu Y, Lei F. 2016. Evolution of body morphology and beak shape revealed by a morphometric analysis of 14 Paridae species. Front. Zool. 13:30.

Shen X-X, Zhou X, Kominek J, Kurtzman CP, Hittinger CT, Rokas A. 2016. Reconstructing the Backbone of the Saccharomycotina Yeast Phylogeny Using Genome-Scale Data. G3 6:3927–3939.

Silvestro D, Schnitzler J, Zizka G. 2011. A Bayesian framework to estimate diversification rates and their variation through time and space. BMC Evol. Biol. 11:311.

Smith SA, Beaulieu JM, Donoghue MJ. 2010. An uncorrelated relaxed-clock analysis suggests an earlier origin for flowering plants. Proc. Natl. Acad. Sci. U.S.A. 107:5897–5902.

Takezaki N. 2018. Global Rate Variation in Bony Vertebrates. Genome Biol. Evol. 10:1803–1815.

Tamura K, Battistuzzi FU, Billing-Ross P, Murillo O, Filipski A, Kumar S. 2012. Estimating divergence times in large molecular phylogenies. Proc. Natl. Acad. Sci. U.S.A. 109:19333–19338.

Tamura K, Tao Q, Kumar S. 2018. Theoretical foundation of the RelTime method for estimating divergence times from variable evolutionary rates. Mol. Biol. Evol. 35:1170–1782.

Thorne JL, Kishino H, Painter IS. 1998. Estimating the rate of evolution of the rate of molecular evolution. Mol. Biol. Evol. 15:1647–1657.

Wickett NJ, Mirarab S, Nguyen N, Warnow T, Carpenter E, Matasci N, Ayyampalayam S, Barker MS, Burleigh JG, Gitzendanner MA, et al. 2014. Phylotranscriptomic analysis of the origin and early diversification of land plants. Proc. Natl. Acad. Sci. U.S.A. 111:E4859–4868.

Wiens JJ, Moen DS. 2008. Missing data and the accuracy of Bayesian phylogenetics. J. Syst. Evol. 46:307–314.

Wikström N, Savolainen V, Chase MW. 2001. Evolution of the angiosperms: calibrating the family tree. Proc. R. Soc. B 268:2211–2220.

Willcock S, Martínez-López J, Hooftman DA, Bagstad KJ, Balbi S, Marzo A, Prato C, Sciandrello S, Signorello G, Voigt B, et al. 2018. Machine learning for ecosystem services. Ecosyst. Serv.

Wyles JS, Kunkel JG, Wilson AC. 1983. Birds, behavior, and anatomical evolution. Proc. Natl. Acad. Sci. U.S.A. 80:4394–4397.

Xi Z, Liu L, Davis CC. 2015. The impact of missing data on species tree estimation. Mol. Biol. Evol. 33:838–860.

Xie W, Lewis PO, Fan Y, Kuo L, Chen M-H. 2011. Improving marginal likelihood estimation for Bayesian phylogenetic model selection. Syst. Biol. 60:150–160.

Yang Z. 2007. PAML 4: phylogenetic analysis by maximum likelihood. Mol. Biol. Evol. 24:1586–1591.

